# A Reference-free Approach for Cell Type Classification with scRNA-seq

**DOI:** 10.1101/2021.05.29.446268

**Authors:** Qi Sun, Yifan Peng, Jinze Liu

**Affiliations:** Department of Computer Science, University of Kentucky, Lexington, KY, 40508, USA; Department of Population Health Sciences, Weill Cornell Medicine, New York, NY 10065, USA; Department of Biostatistics, Virginia Commonwealth University, Richmond, VA 23298, USA

**Keywords:** scRNA-seq, reference-free, alignment-free, simhash, cell type classification

## Abstract

The single-cell RNA sequencing (scRNA-seq) has become a revolutionary technology to detect and characterize distinct cell populations under different biological conditions. Unlike bulk RNA-seq, the expression of genes from scRNA-seq is highly sparse due to limited sequencing depth per cell. This is worsened by tossing away a significant portion of reads that cannot be mapped during gene quantification. To overcome data sparsity and fully utilize original sequences, we propose scSimClassify, a reference-free and alignment-free approach to classify cell types with *k*-mer level features derived from raw reads in a scRNA-seq experiment. The major contribution of scSimClassify is the simhash method compressing *k*-mers with similar abundance profiles into groups. The compressed *k*-mer groups (CKGs) serve as the aggregated *k*-mer level features for cell type classification. We evaluate the performance of CKG features for predicting cell types in four scRNA-seq datasets comparing four state-of-the-art classification methods as well as two scRNA-seq specific algorithms. Our experiments demonstrate that the CKG features lend themselves to better performance than traditional gene expression features in scRNA-seq classification accuracy in the majority of cases. Because CKG features can be efficiently derived from raw reads without a resource-intensive alignment process, scSimClassify offers an efficient alternative to help scientists rapidly classify cell types without relying on reference sequences. The current version of scSimClassify is implemented in python and can be found at https://github.com/digi2002/scSimClassify.

## 1 Introduction

Cataloging cells is crucial for understanding the organization of cells, disease mechanisms, and even treatment respondences. Single-cell RNA sequencing (scRNA-seq) makes it possible to identify cell subpopulations by exploring the unique transcriptomic profile of each cell. Clustering is the most popularly used approach to partition cells based on transcriptome similarity in an unsupervised fashion [1]. However, this requires well-established knowledge of biomarkers for cell type annotation as well as cell populations. Unfortunately, such information is often unavailable prior to the scRNA-seq experiments [2]. Therefore, researchers turn to other machine learning approaches, such as supervised classification, to annotate cells automatically [3].

Recently, Abdelaal *et al*. [3] benchmarked 22 classification methods for scRNA-seq cell type identification. All of these classification approaches utilized gene expression profiles of individual cells as classification features. The study included many conventional classifiers such as support vector machine (SVM) and random forest (RF) in addition to a few recently developed single cell-specific classifiers including ACTINN [4] and scPred [5]. The study demonstrated the efficacy of the gene profile-based approach in cell type identification. In a different study, Arvind Iyer *et al*. [6] classified cell types by naive Bayes, gradient boosting machine, and random forest fitted with gene expression profiles to recognize circulating tumor cells (CTCs) of diverse phenotypes.

However, scRNA-seq data is notorious for its relatively low sequencing depth resulting in highly sparse gene expression across all cells [7]. To make things worse, read alignment to the reference genome often filters out many unmapped reads. It is not uncommon that about half of the reads are thrown out prior to the final analysis [8]. Note that not all unmapped reads are bad reads. Using standard reference genomes may eliminate reads representing significant variations in a particular subject, cell type, or disease genome. Last but not least, aligning read to the reference genome to derive a gene-cell count matrix is typically the most time-consuming step of the process.

To overcome these limitations, we develop a reference-free approach for cell type classification sidestepping read mapping step [9]. Specifically, it explores novel features derived from the entirety of the reads. Instead of using gene expression features derived from scRNA-seq reads, we use *k*-mers, often referred to as the genomic words, as features for classification. Intuitively, these genomic words can be extracted from reads in a scRNA-seq sample. Each of these “words” is associated with its own “frequency” or abundance, which is defined as the number of times that a *k*-mer appears in a sample. The change of gene/transcript expression will correspondingly affect the abundances of *k*-mers identifying them. Thus *k*-mers and their abundances can be used as features for classification due to their strong association with the expression of genes/transcripts. The advantage here is that *k*-mers can be easily derived from reads without alignment to references. In the meantime, the derived *k*-mer set also captures cell and subject-specific variations that do not fit standard reference genomes.

The challenge associated with *k*-mer based features is the huge set of unique *k*-mers, which can be in hundreds of millions depending on sequencing depth. However, a large set of features in the size of hundreds of millions is not a blessing for classification to achieve better accuracy and scalability. We observe that many *k*-mers may be expressed very similarly even across samples, such as a group of *k*-mers unique to the same gene/transcript. These *k*-mers are redundant to each other to represent the true *k*-mer feature space. Clustering is one of the popular unsupervised approaches to group similar objects [10]. Unfortunately, they are not feasible to group abundance profiles of *k*-mers due to the unknown number of clusters as well as high computational cost when dealing with a large amount of *k*-mers directly. Various approaches have been developed in the past in the field of metagenomics classification to reduce the set of *k*-mer features, but they are restricted to applications with only case and control experiments. In this case, *k*-mers that can significantly differentiate case and control were selected for further classification [11,12]. Unfortunately, such approach cannot be easily applied as *k*-mer abundances in scRNA-seq cannot be set up as a two-group comparison. Often times, cell type classification is a multi-class classification problem with half a dozen or more cell types in a single experiment.

In this paper, we propose scSimClassify, a reference-free approach for cell type classification. The scSimClassify reduces the original *k*-mer feature space by partitioning it into subsets of *k*-mers with similar abundance profiles across a variety of cell types via an unsupervised approach. This is achieved by repurposing simhash [13], an extremely fast and effective algorithm that can automatically detect similar items within a large set. We evaluate the performance of scSimClassify on scRNA-seq datasets generated from breast cancer tissues with tumor and immune cell populations, as well as blood samples for studying peripheral blood mononuclear cells (PBMCs) in COVID-19 and influenza patients. Our experiments demonstrate that scSimClassify can accurately identify cell types with the aggregated *k*-mer profiles (CKG features). We also find that the top-ranked CKG features are biologically meaningful in consistency with gene expression features. To the best of our knowledge, scSimClassify is the first reference-free method for multi-class cell type classification based on *k*-mer level information. Besides improving general classification accuracy, our approach also makes it possible to classify cell types with incomplete or even unknown references.

## 2 Methods

Figure 1 describes an overview of scSimClassify training steps for cell type classification. scSimClassify takes the real-value *k*-mer abundance matrix as the input. The matrix is assembled from cells sequenced by scRNA-seq. Here we define *k*-mers sharing similar abundance profiles across cells in a training set as similar *k*-mers. To reduce the size of the input, simhash-based group generator (simGG) is implemented in three steps: (1) generate *k*-mers’ fingerprints, (2) group similar *k*-mers into a compressed *k*-mer group (CKG), and (3) determine CKG abundance matrix. Finally, scSimClassify uses the CKG abundance matrix for cell type classification. Table 1 includes formal notations that will be used in the following sections.

**Fig. 1:**
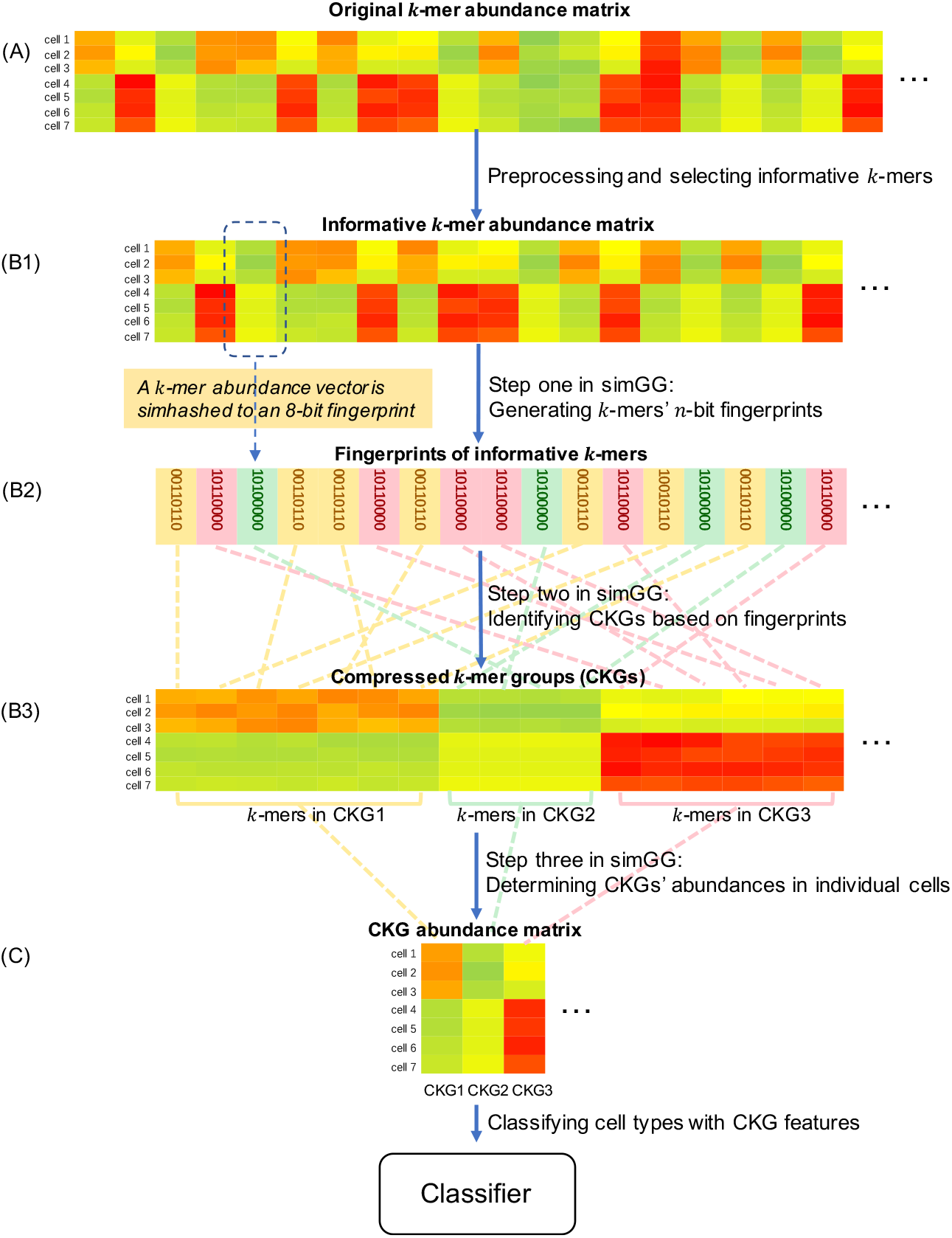
An overview of scSimClassify training steps for cell type classification. (A) The *k*-mers and their abundances in individual cells are obtained as the original input. Preprocessing is applied to the original *k*-mer abundance matrix to filter out noises and systematic variations. Then, informative *k*-mers are selected based on their abundance variability. (B1) In the first step of simGG, *k*-mer abundance vectors are converted to *n*-bit fingerprints through simhash (taking *n* = 8 as an example). (B2) In the second step of simGG, compressed *k*-mer groups (CKGs) are identified based on *k*-mers’ fingerprints. Each CKG contains a set of *k*-mers sharing the same fingerprint. (B3) In the third step of simGG, the abundance of a CKG in a cell is determined by averaging abundances of *k*-mers following the removal of abundance outliers in the same group. (C) Finally, a classifier is trained with the cells represented by CKG features. In this figure, colors of abundance matrices indicate the values of abundances.

**Table 1:** Formal notations in this paper

|  |  |  |  |
| --- | --- | --- | --- |
| $i$ | the index of a $k$-mer | $d$ | the index of a cell |
| $I$ | unique $k$-mers in a training set | $I'$ ($I' \subset I$) | unique informative $k$-mers in a training set |
| $D$ | cells in a training set | $\omega_{di}$ | the abundance of $k$-mer $i$ in cell $d$ |
| $\mathcal{R}_i = \{\omega_{di}\}_{d \in D}$ | a $k$-mer abundance vector, representing a $k$-mer $i$ with abundances across cells in a training set | $\mathcal{C}_d = \{\omega_{di}\}_{i \in I}$ | a cell $d$ represented with an abundance vector across $k$-mers in a training set |
| $g$ | the index of a CKG | $G$ | a set of CKGs |

### 2.1 Preprocessing and selection of informative *k*-mers

The counting of *k*-mer abundances in scRNA-seq reads of individual cells is carried out using jellyfish [14]. Only the canonical form of a *k*-mer sequence is kept, i.e., the lexicographical minimum of itself and its reverse complementary sequence. The original *k*-mer abundance matrix as shown in Figure 1 (A) is further processed based on three principles: (1) normalize *k*-mer abundance within individual cells; (2) filter out *k*-mers with sparse expressions across cells in a training set; (3) select informative *k*-mers with high abundance variations across cells in a training set.

To allow for a fair comparison across cells with variable sequencing depth, we normalize the original *k*-mer abundance of each cell, i.e., 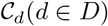, by the total number of sequenced reads in cell *d*.

Often, *k*-mers with sporadic expression across the cell populations may be unreliable due to sequencing errors. We define 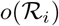 as the occurrence of *k*-mer *i* ∈ *I*, which is the number of nonzero entries in 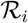. A *k*-mer is removed if it appears in only a small percentage of cells in a training set, i.e., 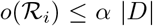. The default setting for *α* is 10%.

Note that not all *k*-mers are equally important for classification purposes. For example, some *k*-mers from housekeeping genes may have very consistent abundances across all cells. Such *k*-mers may not be useful in differentiating cell types. We assume the abundance vector of an informative *k*-mer exhibits a high standard deviation. Let 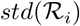 be the standard deviation of *k*-mer abundance vector 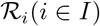. A *k*-mer is selected as an informative *k*-mer if its 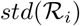 is among the top β% in *k*-mer set *I*. The default setting for *β* is 5. The set of informative *k*-mers *I*’ (*I*’ ⊂ *I*) will be the input of the next step.

### 2.2 Simhash-based group generator (simGG)

As mentioned in Introduction, *k*-mers may originate from the same gene/transcript, sharing similar abundance profiles across cells. Such *k*-mers can be redundant to represent cells. Therefore, we want to group similar *k*-mers into CKGs based on their corresponding abundance vectors across cells in a training set. However, conventional clustering algorithms are not scalable due to the presence of a huge set of *k*-mers. Even *k*-means clustering can not be applied due to the lack of knowledge on the number of clusters.

In this study, we utilize the locality sensitive hashing (LSH) [15], an approximate algorithm that is applicable to objects on a large scale, to detect similar *k*-mers. The underlying idea of LSH is to hash objects with similar features to similar hash values such that object similarity could be determined by comparing their corresponding hash values. Here, we adapt the simhash method [13] to group *k*-mers sharing similar abundance vectors. Simhash was originally developed to identify documents with similar word vectors in a large corpus. The simhash method is one of LSH functions that can represent feature vectors of objects in the continuous space with *n*-bit fingerprints in a binary form. It has the property that the more similar the objects are, the smaller the Hamming distance between their fingerprints, and the higher probability that they share the same fingerprints. Our proposed simhash-based method, named simGG, has the following steps:

**Algorithm 1.**
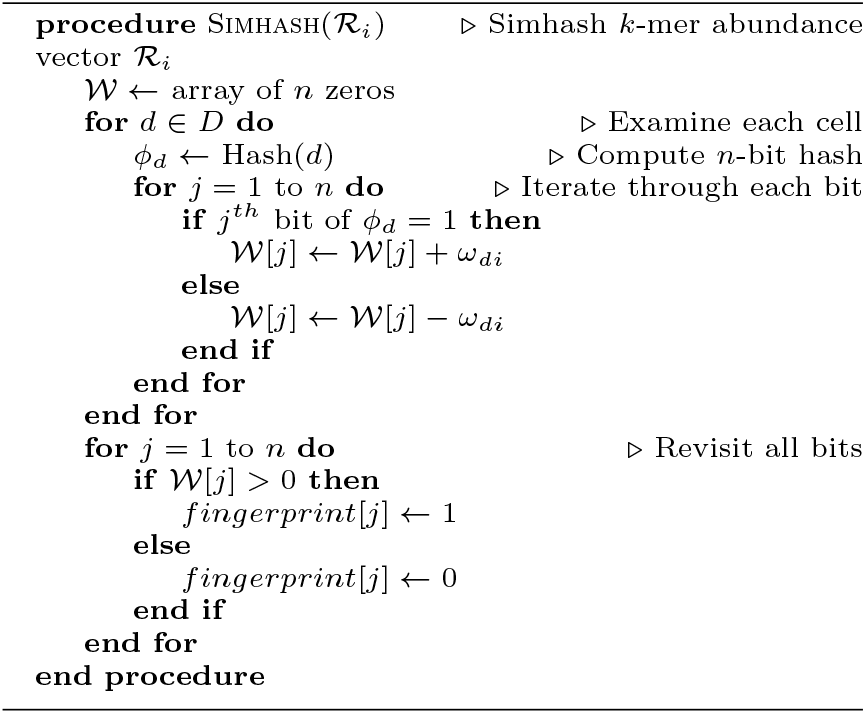
Pseudocode of the simhash algorithm.

#### Generate *k*-mers’ *n*-bit fingerprints

Given a point in space (in this case, a *k*-mer abundance vector), the simhash method generates an *n*-bit fingerprint by determining the point’s relative location among *n* generated hyperplanes. Each bit of the fingerprint corresponds to a hyperplane. The bit’s value is set to 1 if the point is above the corresponding hyperplane; otherwise, it is set to 0. Two points with the same *n*-bit fingerprint indicate that they are very close as none of the *n* hyperplanes is able to separate them. Therefore, using more hyperplanes (larger n) often result in a more accurate similarity estimation for *k*-mer abundance vectors as space is split into much smaller regions.

To speed up the performance and avoid storing hyperplanes, we implement the simhash method as the pseudocode given in Algorithm 1 [16]. The steps to map a *k*-mer abundance vector 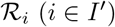 to a *n*-bit fingerprint start by initializing a temporary array 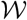 with *n* zeros. Next, the algorithm generates an *n*-bit hash *ϕ_d_* for each cell *d* in 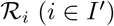 with a consistent hashing mechanism md5 [17]. For each bit of *ϕ_d_*, it decides to add or subtract *ω_di_*, the abundance of *k*-mer *i* in cell *d*, to/from 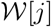 based on whether the *j*-th bit of *ϕ_d_* is one or zero. After all the cells of 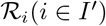 are processed, *j^th^* bit in fingerprint is obtained by setting 1 if 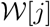 is positive, otherwise setting to 0. Therefore, a *k*-mer abundance vector 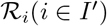 is mapped to [0,2^*n*^] *n*-bit fingerprint values as shown in Figure 1 (B1) and (B2).

#### Identify compressed *k*-mer groups (CKGs)

Based on the property of simhash, two *k*-mers are considered similar if the Hamming distance between their corresponding fingerprints is very small. Considering the large scale of *k*-mers and similarity identification performance, we use the strictest measure to identify similar *k*-mers by checking if Hamming distance is zero or not [18]. Thereby we do not need to compute Hamming distances of all pairs of *k*-mers’ fingerprints. We define a CKG as a group of similar *k*-mers if they share exactly the same fingerprint. An example of CKGs is provided in Figure 1 (B3).

To group similar *k*-mers, a naive clustering method takes *O*(|*I*’|^2^|*D*|) time. In comparison, the complexity of simGG is bounded by *O*(ļ*I*’ļ log ļ*I*’ļ). The algorithm first simhashes informative *k*-mer abundance vectors 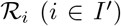 to *n*-bit fingerprints with *O*(|*I*’|) time complexity. This is followed by the identification of the *k*-mers with the same fingerprints through sorting with *O*(|*I*’| log |*I*’|) time complexity. Additionally, both of the steps in simGG can be executed in parallel computing [19], which may further reduces the running time.

#### Determine CKGs’ abundances in individual cells

In this step, the pre-built CKGs from a training set are used to aggregate *k*-mer abundances into CKGs’ abundances for both training and test sets. In general, the abundances of *k*-mers belonging to the same CKG are similar in an individual cell, as shown in Figure 1 (B3). As a result, we can compress those *k*-mers’ abundances into a single abundance to represent the expression of a CKG in a cell. Given *k*-mers belonging to cell d and CKG *g*, we first filter out the outliers whose abundances fall outside of two standard deviations from the mean abundance and then average the abundances of the remaining *k*-mers as the abundance of CKG g for cell d. We iteratively determine the abundance of each CKG for each cell such that each of the cells can be characterized by a set of CKG features. As shown in Figure 1 (A) and (C), the feature size of individual cells is reduced from the original *k*-mer size |*I*| to CKG feature size |*G*|.

### 2.3 Classification Algorithms

During the classification process, we adopt a variety of classification algorithms [3, 6] to classify cell types with CKG features. These methods include random forest (RF) [20], gradient boosting machine (GBM) [21], multilayer perceptron (MLP) [22] and support vector machine (SVM) [23]. The four classifiers are selected to classify cell types with CKG features as they represent four branches of the general classification algorithms. RF and GBM are tree-based ensemble methods that randomly consider a subset of features to build the classifier. The difference between them is that RF builds trees independently, while GBM builds one tree at a time to correct decision trees that come before it. MLP is a kind of artificial neural networks that considers all the features to determine the data classes. SVM finds a plane with the maximum margin to separate two classes of data points. Benchmarking on these classifiers allows us to investigate how different CKG features perform on each type of the state-of-the-art classifiers.

## 3 Experimentation and evaluation

The goal of our experiment was to evaluate scSimClassify for cell type classification using scRNA-seq data. Here we compared the performance of CKG features and commonly used gene expression features in the application of cell type classification. We conducted thorough comparisons among numerous general purpose classifiers (RF, GBM, MLP and SVM) between the two types of features. Benchmarking gene expression based classification methods to automatically assign cell identities, Abdelaal *et al*. [3] concluded that ACTINN [4] and scPred [5] performed well on most datasets as single cell-specific classifiers. Therefore, we also compared classification performance with ACTINN and scPred based on gene expression features.

### 3.1 Experiment configuration

#### Datasets

We identified four datasets (Chuang [24], Karaayvaz [25], PBMC3k [26] and Lee [27]) for evaluation in our experiments. They include two datasets of similar cell types in breast cancer tissues, and two datasets of peripheral blood mononuclear cells (PBMCs) (Table A.1). They vary in the number of cells, cell populations, and sequencing protocols.

Both Chuang’s and Karaayvaz’s datasets were sequenced from breast cancer tissues using Smartseq-2 technology where full-length transcripts were sequenced within individual cells. Their associated experiments aimed at revealing the characteristics of breast cancer subtypes shaped by tumor cells and immune cells in the microenvironment [24, 25]. The Chuang’s dataset with GEO accession number GSE75688 [24] contains 317 epithelial breast cancer cells, 175 immune cells, and 23 stromal cells. The epithelial breast cancer cells are further divided into four subpopulations: 73 luminal A subtypes, 25 luminal B subtypes, 130 HER2 subtypes, and 89 triple-negative breast cancer (TNBC) subtypes. And 175 immune cells can be further classified into three categories: 83 B cells, 54 T cells, and 38 macrophages. In all, it consists of eight types of cells. The Karaayvaz’s dataset with GEO accession number GSE118389 [25] contains 1098 cells originated from five different cell type populations: 868 epithelial breast cancer cells, 94 stromal cells, 64 macrophages, 53 T cells, and 19 B cells. Both datasets share five common cell types: epithelial breast cancer cells, stromal cells, B cells, T cells, and macrophages, making it possible to classify cell types across datasets.

The PBMC3k and Lee’s datasets are human PBMC datasets. They were sequenced by 10x genomics, which only sequenced the 3’-end of the transcripts and generate a relatively low number of reads. The scRNAseq data and its gene expression profiles from PBMC3k dataset are freely available from 10X Genomics [26] with nine identified cell types. This is a well-analyzed dataset with the ground truth cell type assigned by Seurat clustering protocol [28]. Lee’s study performed scRNA-seq using PBMCs to identify factors associated with the development of severe COVID-19 infection. We randomly selected three samples from Lee’s dataset with GEO accession number GSE149689. They were PBMCs from a healthy donor (HD), a patient with severe influenza (FLU), and a patient with COVID-19. Also, six shared cell types between PBMC3k and Lee’s dataset made cross-dataset classification possible.

#### Features

Gene expression data associated with the original scRNA-seq data from each dataset was downloaded from the GEO repository or 10X Genomics. We followed QC criteria as used in their original studies [24,27] to discard low-expressed or unexpressed genes. We referred to “gene expression features” as GE in the following sections.

We generate several variations of CKG features as illustrated in Methods with different combinations of *k*-mer length, k and simhash fingerprint size, n. The *k*-mer length is either 16 or 21 [29], while fingerprint bit size for simhash can be either 16-bit or 32-bit. To investigate whether the reference genome is essential for the cell type classification, we generated two categories of *k*-mers as inputs, as listed in Table 2. The first category (on the left in Table 2) was generated without reference-based selection, containing *k*-mers derived from all the reads in scRNA-seq data; The second (on the right in Table 2) contained only *k*-mers derived from reads that can be mapped to the reference genome. To obtain mapped reads in Chuang’s dataset, the scRNA-seq reads were aligned to human genome reference sequences (hg19) using the 2-pass mode of STAR (default parameters) [30], following the same alignment procedure for gene expression quantification [24]. As for PBMC3k dataset, we obtained read alignment information from the downloaded BAM file.

**Table 2:** The nomenclature (id) of CKG feature variations with different combination of the parameter values. The variations on the left are reference-free and take all *k*-mers to generate CKG features. The variations on the right only take *k*-mers that can be mapped to the reference genome to generate CKG features.

| id | $k$ | $n$ | read type | id | $k$ | $n$ | read type |
| --- | --- | --- | --- | --- | --- | --- | --- |
| allk21n16 | 21 | 16 | all | mappedk21n16 | 21 | 16 | mapped |
| allk21n32 | 21 | 32 | all | mappedk21n32 | 21 | 32 | mapped |
| allk16n16 | 16 | 16 | all | mappedk16n16 | 16 | 16 | mapped |
| allk16n32 | 16 | 32 | all | mappedk16n32 | 16 | 32 | mapped |

Due to different sequencing protocols, the average reads per cell on PBMC3k is around 69,000 reads per cell in comparison to over 10 million reads per cell in Chuang’s dataset. Therefore we set the *k*-mer filtering threshold described in section 2.1 to be *α* =0.5% to retain sufficient *k*-mers to generate CKG features in the PBMC3k dataset, comparing to the default setting of *α* =10% as in Chuang’s dataset.

#### Classifier configuration and performance metrics

We applied MLP, RF, GBM, and SVM to classify cells with each feature set. A grid search to identify optimal hyperparameter combination was performed for all classifiers. The hyperparameter searching space for RF and GBM was the maximum tree depth (2/6/10), number of estimators (10/50/100), and a maximum of features to look for best split (“sqrt” / “log” / “None”). SVM selected parameters from the type of kernel (linear/rbf) and the margin error controller (0.0001/0.001/0.01). The options for MLP were the number of the hidden layer (1/2) and the dropout rate (0.4/0.5/0.6). The number of neurons in each layer was the average of neuron numbers of its previous layer and output layer. RF, GBM and SVM were implemented via the scikit-learn library [31], and the MLP was implemented in Keras [32]. We run ACTINN and scPred with their defaulting settings after downloading scripts or installing packages from their respective websites.

To evaluate the performance of multi-class classification on imbalanced data, we calculated accuracy, F1 score by the module in the scikit-learn library. Each class provided a weighted contribution to F1 score [31].

### 3.2 Evaluation of Intra-Dataset cell type classification

In this experiment, we evaluated the scSimClassify’s performance by training and testing subsets of cells included in the same scRNA-seq data. We named this an Intra-Dataset evaluation. The comparisons were made by reporting results from the following groups: (a) scSimClassify with general purpose classifier MLP, RF, GBM and SVM. The features were GE, and 8 variations of CKGs (Table 2). (b) ACTINN, scPred with GE feature set. The stratified 5-fold cross-validation was used to select the best hyperparameter combination for each classifier and feature set in scSimClassify. For all pipelines, five independent repetitions of 5-fold cross-validation were performed to determine the classification results.

#### Comparison between GE and CKG features

We used two datasets, the Chuang’s dataset as well as PBMC3k dataset, to compare the performance of GE and CKE features in their ability for cell type identification. As reported in Table 3, the overall winner for Chuang’s dataset is CKG feature classified by MLP, and CKG feature classified by SVM wins the best classification performance on PBMC3k dataset. All the general purpose classifiers in scSimClassify outperform scRNA-seq specific classifier ACTINN and scPred trained with gene expression features. For each general classification model, using CKG features quite consistently improves the overall classification accuracy over GE features. This supports our hypothesis that *k*-mer level features without gene annotation are sufficient for cell classification.

**Table 3:**
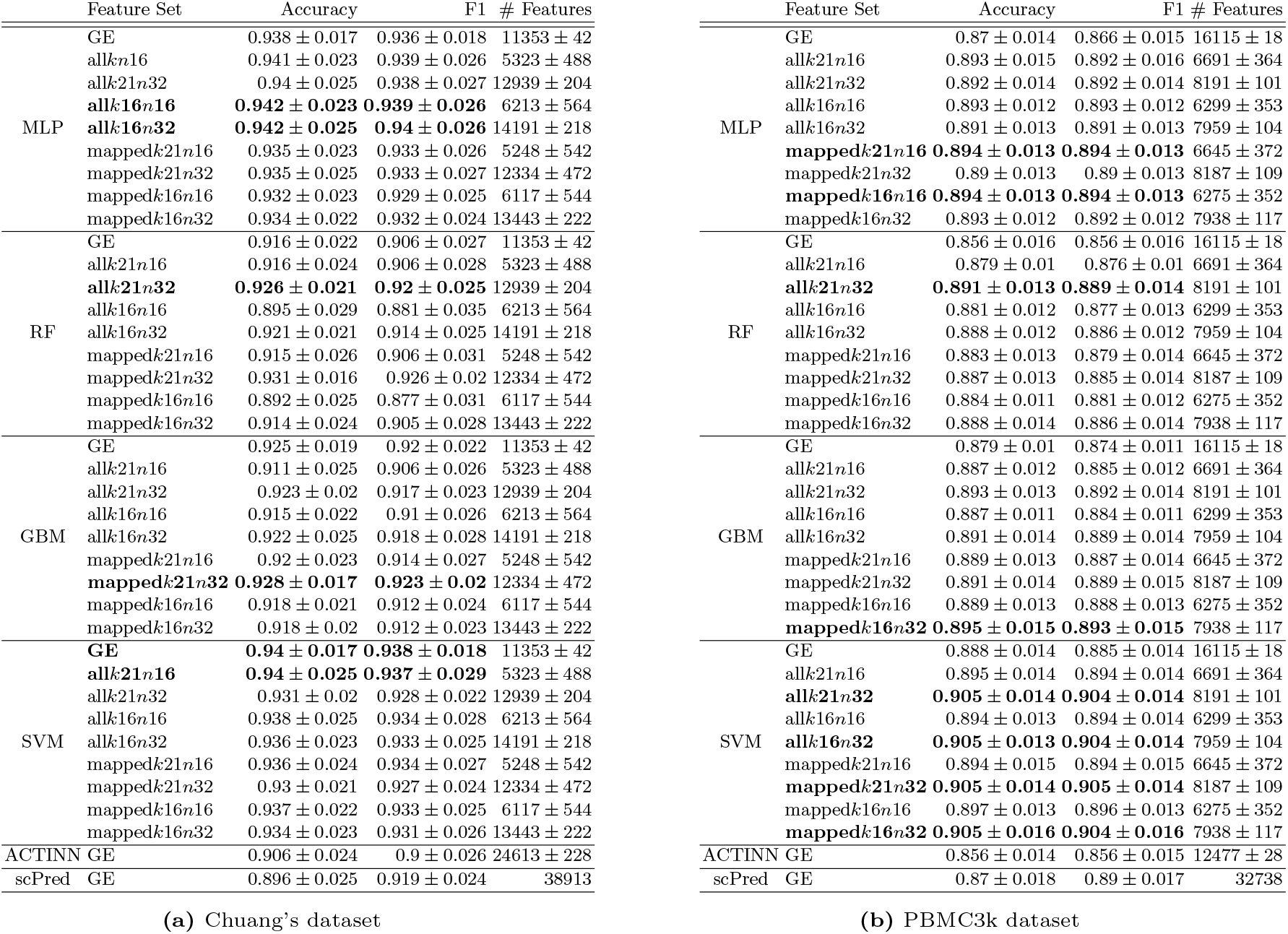
Comparison of Intra-Dataset cell type classification performance among scSimClassify using GE features and 8 variations of CKG features (listed in Table 2), as well as ACTINN and scPred with GE features. The mean and standard deviation are recorded for different evaluation metrics after five repetitions of 5-fold cross-validation. The best performances in each classifier are highlighted in bold.

#### Evaluation of CKG feature variations

In this experiment, we also conducted thorough comparisons of the 8 variations of CKG features to understand the effect of parameters *n, k*, and read types on the CKG performance.

##### CKG feature variations with different values of k

By fixing the size of fingerprints, classifiers, and read types, 21-mer CKG feature variations represent the same or better performance comparing with 16-mer CKG feature variations in 10 cases out of 16 comparisons on both datasets. For example, the performance of allk21n16 is 2.1% better than allk16n16 for RF in accuracy in Chuang’s dataset. It indicates that more unique *k*-mers lead to a finer resolution in representing gene diversity. This can ultimately result in better classification performance.

##### CKG feature variations with different values of n

While we fixed *k*-mer length, classifiers, and read types, the performances of CKG feature variations grouped by 32-bit fingerprints are better than 16-bit fingerprints in around two-thirds of 16 comparisons on both datasets. Theoretically, using the 32-bit fingerprints will generate more random hyperplanes to separate the original *k*-mer space, thus creating a more precise categorization of *k*-mer groups than 16-bit fingerprints. It eventually leads to more descriptive CKG features and better classification performances.

##### CKG feature variations with different read types

CKGs derived from *k*-mers of all reads outperform those from mapped reads in three-quarters of comparisons on Chuang’s dataset and half of the comparisons on PBMC3k dataset. This suggests our reference-free approach is able to capture cell type relevant features for classification without preselecting *k*-mers from mapped reads. The *k*-mers from unmapped reads may contribute to the additional performance gain of our reference-free approach.

Based on the performance comparison of variations of CKG features, we selected all*k*21*n*32 as CKG feature set in the following experiments.

#### Evaluation of highly variable features

Inferring highly variable features is a common step in current bioinformatics analysis [33]. To evaluate the necessity of highly variable features in this study, we selected the top 2000 variable features with default settings of Seurat VST [28] for both GE and CKG (all*k*21*n*32) feature sets. Classification performance on Intra-Datasets with highly variable features is shown in the Appendix Table A.2. Comparing classification performance based on highly variable features and all features (Table 2,Table A.2), there is no clear winner for cell type classification from both datasets and both feature sets. The comparison results are classifier-dependent and dataset-dependent. Moreover, inferring a subset of features may exclude discriminant sources of variation across cells [5] and introduce feature selection parameters. Therefore we used all the features to classify cell types in this study.

### 3.3 Evaluation of Inter-Dataset cell type classification

In this experiment, we evaluated if scSimClassify trained with one scRNA-seq dataset may be applied to classify cell types in the other, which we referred to as Inter-Dataset classification.

We conducted two sets of Inter-Dataset experiments to predict shared cell types, as mentioned in section 3.1. In one experiment, Chuang’s dataset was used as the training data and the trained model was applied to predict the cell types in Karaayvazr’s dataset. In the other experiment, cell types of three PBMC sets, which were cells from a COVID-19 patient, a FLU patient, and a healthy donor in Lee’s dataset, were identified based on the model trained on PBMC3k dataset. To obtain optimal hyperparameters for the target distribution, we randomly chose 20% and 80% of cells in the targeting sets for validation and testing respectively. The validation set was used to grid search optimal hyperparameter combination [34]. The GE and well-performing CKG feature set, all*k*21*n*32, suggested by Intra-Dataset, were used for Inter-Dataset classification.

Table 4 represents the performance of Inter-Dataset classification between Chuang’s dataset and Karaayvazr’s dataset with averaged results over 5 repetitions. Overall, SVM using CKGs(all*k*21*n*32) shows the highest accuracy for detecting cell types, followed by GBM and ACTINN with GE features. Again the CKG features show competitive performance to GE features in almost all metrics. For MLP, RF and GBM, the CKG feature set (all*k*21*n*32) consistently outperforms GE features in all metrics. The scPred method failed to identify cells in this task even tuning the default parameters. Here Inter-Dataset experiment shows a more pronounced performance gain using CKG features over GE features when compared to the performance on Intra-Dataset experiment of the same configuration.

**Table 4:** Comparison of Inter-Dataset cell type classification performance among scSimClassify using GE and CKG (all*k*21*n*3) feature sets, as well as ACTINN with GE features. The classification models are trained on Chuang’s dataset and tested on Karaayvazr’s dataset. The mean and standard deviation are recorded for different evaluation metrics after five repetitions. The best performance in each classifier is highlighted in bold.

|  | Feature set | Accuracy |  | F1 # Features |
| --- | --- | --- | --- | --- |
| MLP | GE | $0.69 \pm 0.045$ | $0.69 \pm 0.045$ | 11381 |
|  | **$\text{all}k21n32$** | **$0.764 \pm 0.039$** | **$0.764 \pm 0.039$** | 12958 |
| RF | GE | $0.803 \pm 0.007$ | $0.803 \pm 0.007$ | 11381 |
|  | **$\text{all}k21n32$** | **$0.828 \pm 0.003$** | **$0.828 \pm 0.003$** | 12958 |
| GBM | **GE** | **$0.842 \pm 0.005$** | **$0.842 \pm 0.005$** | 11381 |
| | $\text{all}k21n32$ | $0.828 \pm 0.002$ | $0.828 \pm 0.002$ | 12958 |
| SVM | GE | $0.692 \pm 0.007$ | $0.692 \pm 0.007$ | 11381 |
|  | **$\text{all}k21n32$** | **$0.872 \pm 0.003$** | **$0.872 \pm 0.003$** | 12958 |
| ACTINN GE | | $0.838 \pm 0.028$ | $0.852 \pm 0.023$ | 17061 ± 36 |

For PBMC Inter-Dataset classification (Figure 2), there is no winner feature set based on the results from three samples in Lee’s dataset. However, for RF, using CKG features consistently improves the accuracy in comparison to using GE features. As for GBM, CKG features show relatively equivalent performance to GE features. As for MLP and SVM, CKG features outperform GE features in FLU sample.

**Fig. 2:**
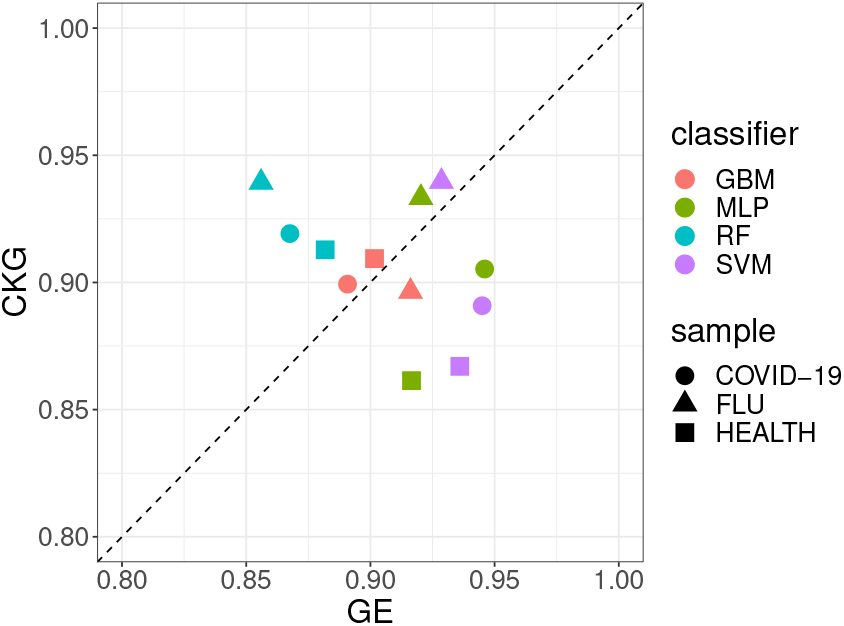
Comparison of CKG features (all*k*21*n*32) and GE features for Inter-Dataset PBMC classification. Each point in the scatter plot shows the accuracy using CKGs vs using GE. Three PBMC samples in Lee’s dataset are used. Each cell in the sample is classified by four classifiers.

#### 3.4 Feature interpretation

In this section, we tried to identify the biological origin of important CKG features and assessed whether they were biologically meaningful.

A CKG was formed by *k*-mers sharing similar abundance profiles across cells. These *k*-mers might be from the same gene, genes sharing significant sequence similarity (such as gene families), or even co-regulated genes. Here we defined that a CKG as a single-gene CKG if more than 90% of *k*-mers in it can be mapped to one and only one gene. For those CKGs from genes with shared subsequences or potential co-regulated genes, we defined them as multi-gene CKGs if at least 45% of *k*-mers in them can be mapped to each gene. Except for single-gene and multi-gene CKGs, we categorized the remaining CKGs in the CKG feature sets as unannotated CKGs. To identify the annotation for a CKG, we ran a blast search to determine each *k*-mers’ gene association against protein-coding reference transcriptome (hg19).

We generated CKG feature sets (all*k*21*n*32) from Chuang’s and PBMC3k datasets respectively to investigate CKG annotation distribution. Ranking CKG feature importance by trained tree-based models (RF, GBM), we analyzed annotation distributions of top N of the most important CKGs in the feature sets by changing the value of N (Figure 3, Figure A.1). Setting N as the number of CKGs in a feature set (the last stacked bar in Figure 3), it shows that 67% of CKGs are single-gene CKGs, 5% of them are multi-gene CKGs among feature set generated from Chuang’s dataset, while the corresponding proportions for PBMC3k dataset are 80.7% and 7.1%. It supports that simGG is capable of statistically grouping *k*-mers from a gene or multiple genes. Most of the genes associated with multi-gene CKGs come from the same gene families sharing subsequences. As expected, the proportion of single-gene CKG increased while decreasing N and selecting a relatively small set of the most important CKG features. However, there still exist multi-gene and unannotated CKGs even when N is as small as 50. This indicates that both multi-gene and unannotated CKGs carry differentiate information for cell type classification.

**Fig. 3:**
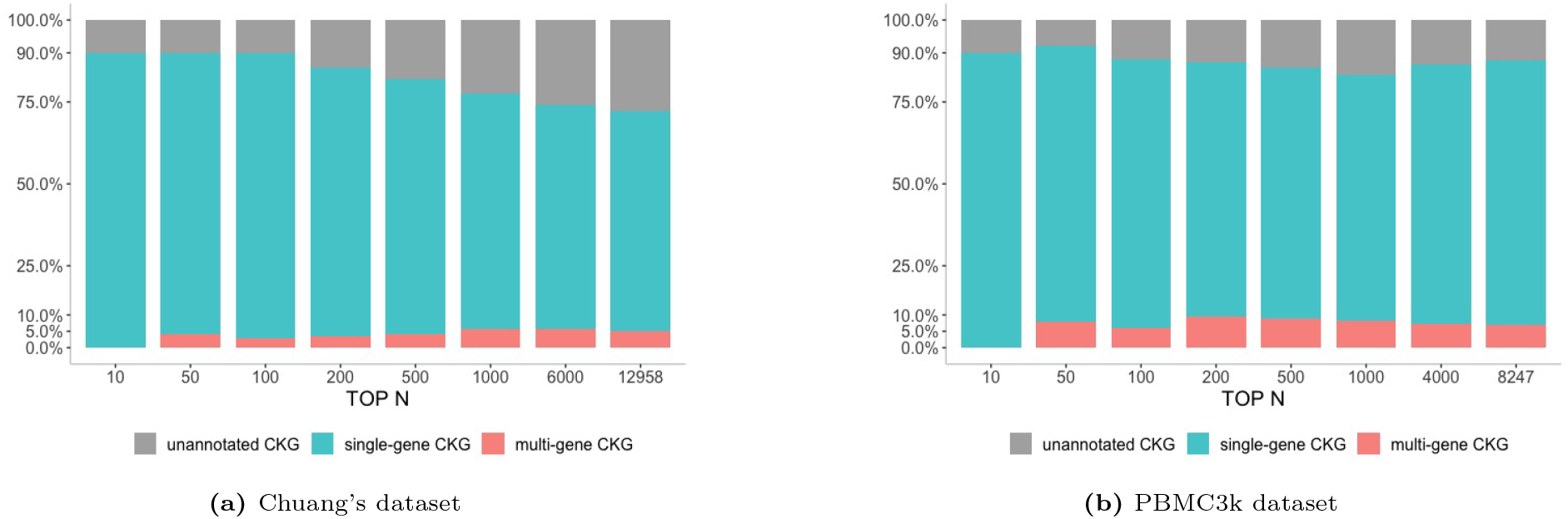
Distributions of three categories of CKGs in terms of their association with known gene annotation among top N of the most important CKGs derived from RF models. The CKG feature sets (all*k*21*n*32) are generated by Chuang’s dataset (a) and PBMC3k dataset (b) respectively. The last stacked bar shows the category distribution of the whole CKG feature set.

We next analyzed the common genes shared within CKGs’ gene annotation and GE. Here we focused on exploring features with a significant contribution to the classification. The top 10 most important GE and CKG features were derived from tree-based models used in Intra-Dataset experiments. From five repetitions of 5-fold cross-validation on Intra-Dataset, we obtained 25 sets of classification models. For the top 10 most important GE features, its gene set consists of unique genes over 250 genes. For the top 10 most important CKGs (all*k*21*n*32), the gene set consists of unique genes over gene annotations of 250 CKGs.

Given a large proportion of common genes associated with the top 10 most important features in GE and CKGs derived from both RF and GBM (Table 5), we have the following observations. First, a large proportion of these genes are marker genes for each cell type classification task. For Chuang’s dataset, numerous genes, such as PPP1R1B [35], FABP7 [36] and ERBB2 [37], were reported by prior literature showing close associations with breast cancers [35–39]. For PBMC3k dataset, a set of common genes (CCL5, CD14, CD3D, CD79A, CFD, CST3, GNLY, LST1, LYZ, NKG7, S100A4, S100A8, S100A9, TCL1A) were identified as marker genes by Seurat scRNA-seq analysis pipeline [28]. Secondly, the vast majority of common genes, as highlighted in bold, are shared by both RF and GBM classification models. This suggests that common genes from CKG gene annotations and GE can be consistently derived from tree-based models.

**Table 5:**
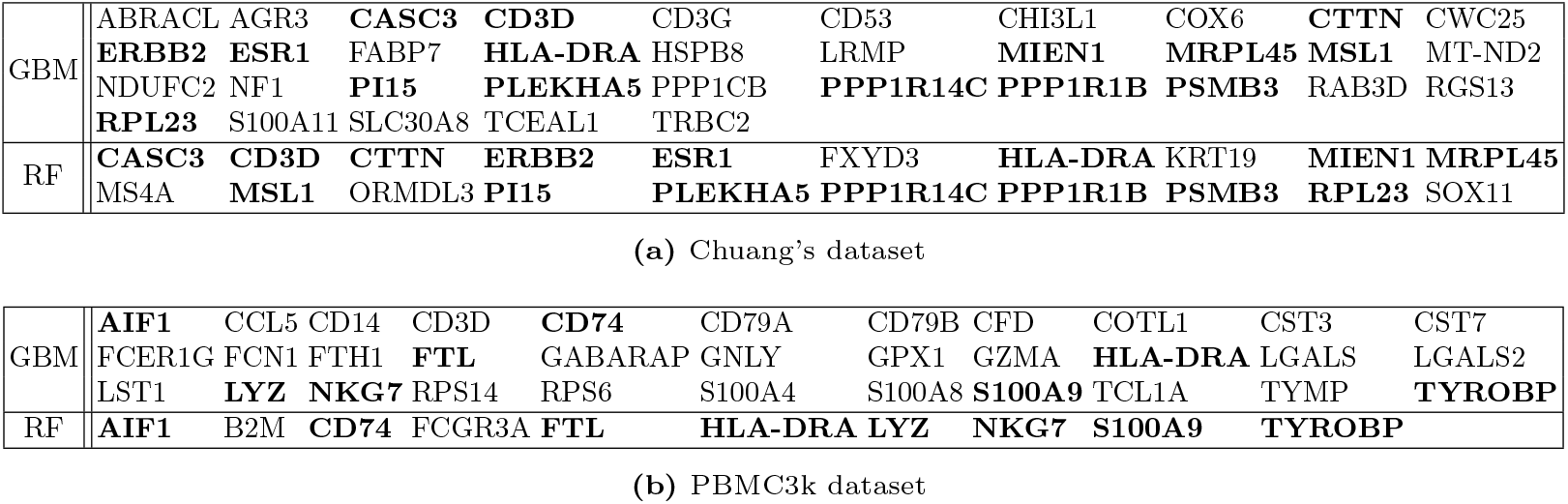
Common genes identified as the top 10 most important genes selected from GE features and associated with the top 10 most important CKG features (all*k*21*n*32) in Intra-Dataset experiments under RF and GBM. The overlaps of common genes between RF and GBM are highlighted in bold.

## 4 Discussion and Conclusion

This paper presents a reference-free classification method for cell type identification in scRNA-seq data. Our method leverages *k*-mer level features from the entirety of the reads for cell type classification without requiring the alignment of reads. This enables the utilization of full sequencing reads especially when the reference genome is unavailable or when the subject genome is highly mutated.

Our experiments on four datasets demonstrate that our proposed CKG features serve as competitive features to gene expression features for cell type classification, which are exhibited across a variety of classification models. This suggests that CKG features can be an effective alternative to gene expression features for cell type identification and can potentially be used in replacement of gene expression features.

In this study, we attempt to interpret CKGs using the *k*-mers associated with genes. We find that our method naturally groups *k*-mers originated from the same gene together. This allows us to annotate CKG features with known genes to assess their biological significance. The significant overlap of gene annotations of top-ranked CKG features with top-ranked genes from GE indicates our method is biologically meaningful. While we demonstrate that CKGs without specific gene annotations are also discriminative for cell types, their potential biological association with mutations and intergenic elements deserves further investigation in future work.

Our future work will focus on three directions: (1) We plan to expand the current evaluation to include more scRNA-seq datasets for validation and benchmarking; (2) We will continue our effort in the biological interpretation of CKG features; (3) We will further optimize configuration parameters such as exploring even larger fingerprint size *n* to see if the performance gain will continue to improve or will plateau at a certain point.

## 5 Acknowledgment

The work was supported by the National Library of Medicine/National Institutes of Health Grant (No 4R00LM013001).

## A Appendix

### A.1 Supplementary Tables

**Table A.1:**
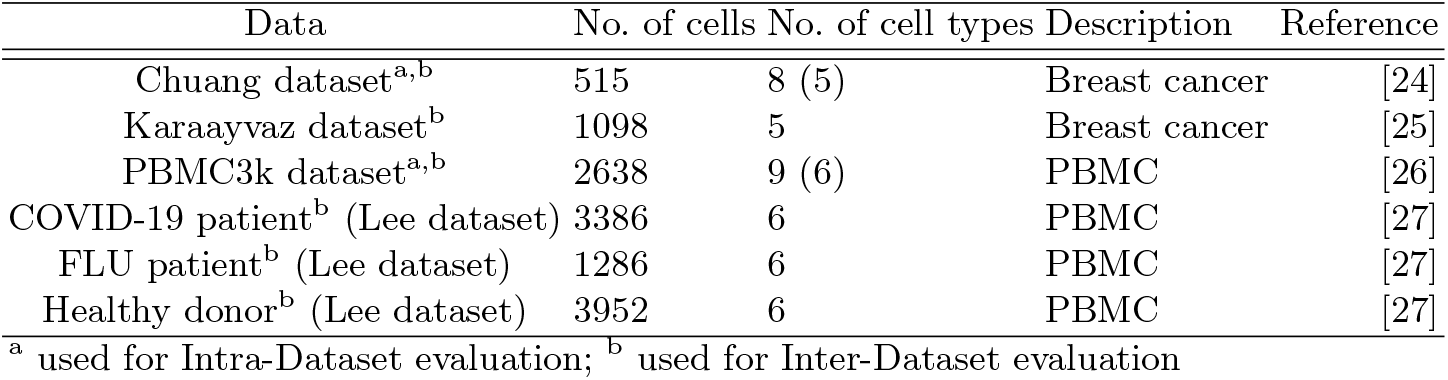
Overview of the datasets used during this study

**Table A.2:**
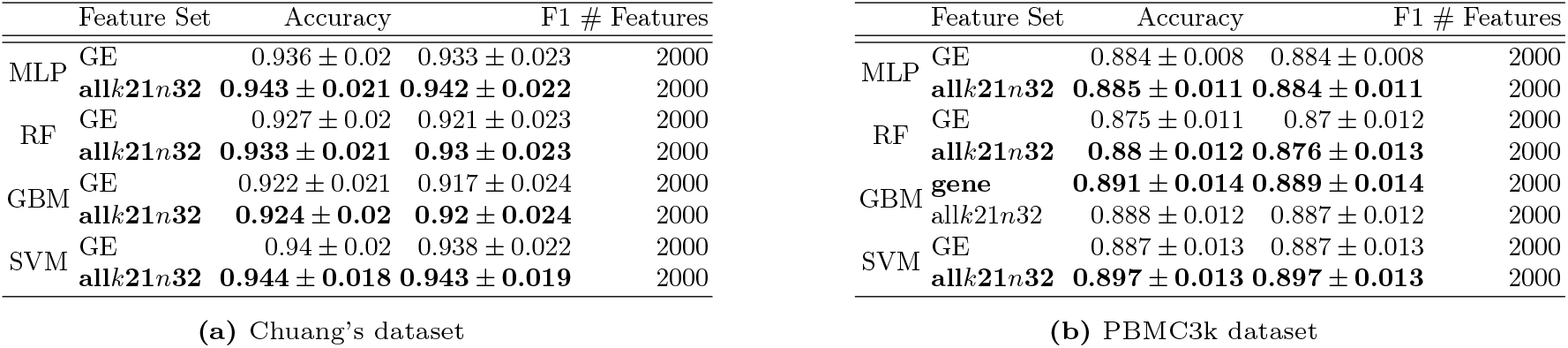
Comparison of Intra-Dataset cell type classification performance among scSimClassify using GE and CKG (all*k*21*n*32) features. Top 2000 highly variable features are selected for training the models. The mean and standard deviation are recorded for different evaluation metrics after five repetitions of 5-fold cross-validation. The best performances in each classifier are highlighted in bold. CKG (all*k*21*n*32) features perform better than GE features for cell type classification in a majority of cases.

### A.2 Supplementary Figure

**Fig. A.1:**
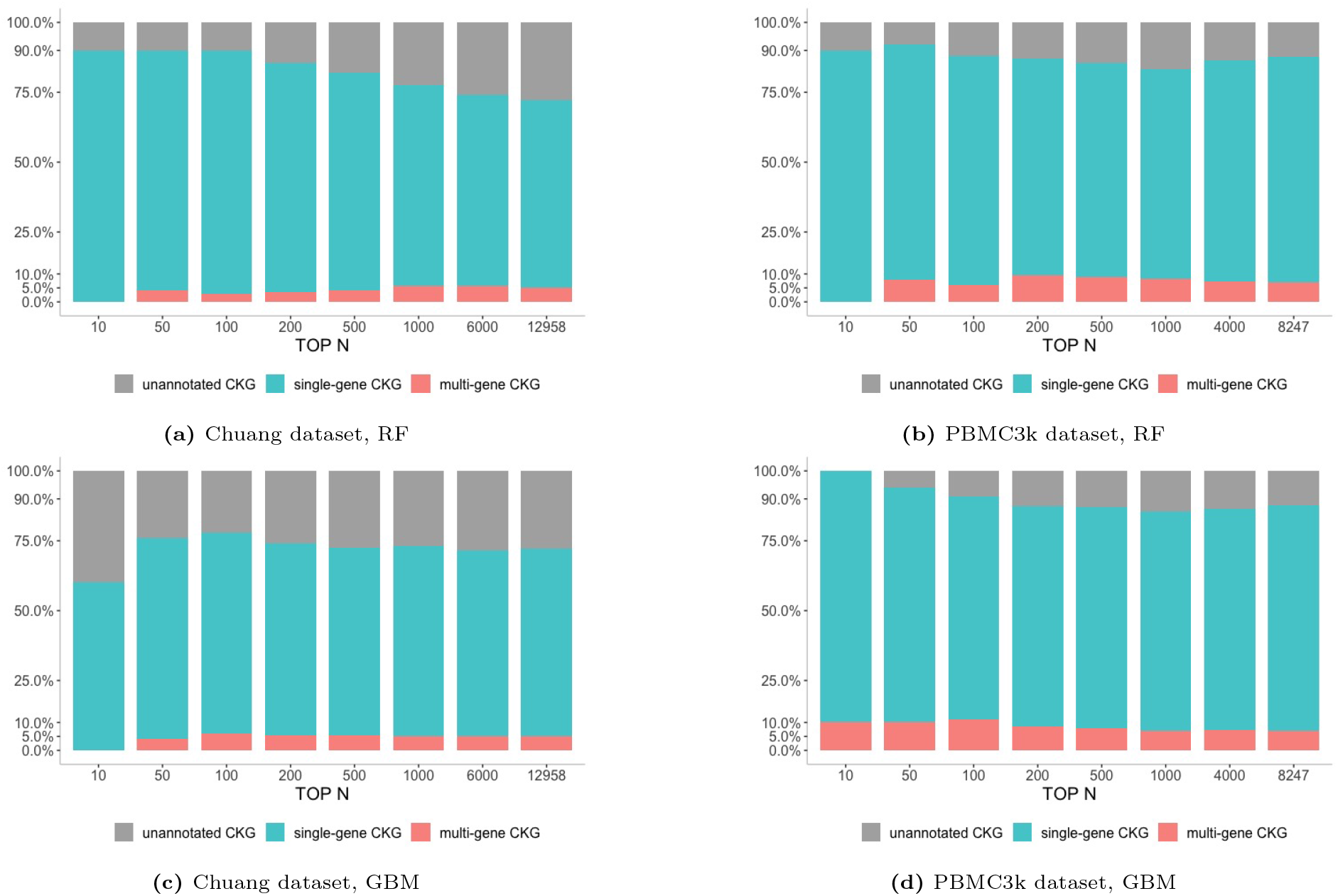
Distributions of three categories of CKGs in terms of their association with known gene annotation among top N of the most important CKGs derived from RF and GBM models. The CKG feature sets (all*k*21*n*32) are generated by Chuang’s dataset (a)(c) and PBMC3k dataset (b)(d) respectively. The last stacked bar shows category distribution of the whole CKG feature set. (The dataset and classifiers are shown below each bar plot).

